# Temporal dynamics and acquisition of Shiga toxin subtype *stx2a* within Shiga toxin-producing *Escherichia coli* in England, 2016 – 2024

**DOI:** 10.64898/2026.04.09.717390

**Authors:** Eleanor H. Hayles, Ella V. Rodwell, David R. Greig, Claire Jenkins, Gemma C. Langridge

**Affiliations:** Quadram Institute Bioscience, Norwich Research Park, Norwich, UK; Norwich Medical School, University of East Anglia, Norwich Research Park, Norwich, UK; Centre for Microbial Interactions, Norwich Research Park, Norwich, UK; Gastrointestinal Bacteria Reference Unit, UK Health Security Agency, London, UK

**Keywords:** Shiga toxin producing *Escherichia coli* (STEC), Shiga toxin, *stx2a*, acquisition, O157, non-O157

## Abstract

Shiga toxin-producing *Escherichia coli* (STEC) are an important public health concern due to their association with foodborne gastroenteritis and severe outcomes including haemolytic uraemic syndrome (HUS), particularly linked to the *stx2a* subtype of the Shiga toxin. We investigated the temporal dynamics and acquisition of *stx2a* among STEC isolates submitted to the United Kingdom Health Security Agency (UKHSA) between 2016 and 2024. 12,888 whole genome STEC sequences and associated metadata were analysed. 31.9% of STEC isolates harboured *stx2a*, spanning 78 O serogroups with a marked shift from STEC O157 to non-O157 serogroups over time. STEC O26:H11 and STEC O145:H28 were the primary drivers of observed increases, most commonly associated with *stx2a* alone or in combination with *stx1a*. The widespread and increasing presence of *stx2a* across the STEC population in England highlights an emerging public health risk and demonstrates the value of routine genomic surveillance in monitoring high-severity Shiga toxin subtypes.

## Introduction

Shiga toxin-producing *Escherichia coli* (STEC) are a diverse group of foodborne gastroenteric pathogens which pose a significant challenge to public health in the England (1). STEC are zoonotic pathogens, with ruminants such as cattle and sheep their main reservoir (2,3). Transmission can occur through the food chain at multiple stages, including pre-harvest through water run-off contaminated with animal faeces and during post-harvest food processing and packaging (4). In the UK, STEC outbreaks from food have been associated with consumption of undercooked meat, unpasteurised dairy products and unwashed vegetables (5–9). Further transmission can occur from direct contact with infected humans or animals (10,11). Whilst reported year-round, seasonality is often observed, with higher cases reported in the late summer months (12–15).

The hallmark of STEC is presence of the biologically potent Shiga toxin (Stx) (16). The *stx* gene is phage encoded, and STEC are capable of being infected with multiple *stx* phages, leading to the potential for multiple *stx* gene presence and expression by a single bacterial cell (17,18). There are two main biological divisions of Stx – Stx1 and Stx2 – which share a 56% similarity at the amino acid level (19,20). Stx1 divides into four further subtypes – *stx1a, stx1c, stx1d and stx1e* - and Stx2 divides into at least 15 subtypes, *stx2a – stx2o* (21,22).

STEC O157 is the most common serogroup associated with human disease, however disease caused by serogroups other than O157 (non-O157 serogroups) have seen a significant rise in recent years, partially owing to a shift from diagnosis by culture-based methods to PCR-based identification of the *stx* gene (23,24). Over 100 non-O157 serogroups are known to have caused serious illness (25). However, the most frequently reported globally are characterised as the “Big Six” non-O157 STEC serogroups – O26, O45, O103, O111, O121 and O145 (26). Of these, O26 and O145 are significantly reported in England each year, alongside O146, O91 and O128 (12–15).

Clinical manifestation of STEC usually presents as gastroenteritis including stomach cramps, fever and bloody diarrhoea (27). More serious conditions can develop in severe cases, including haemolytic uraemic syndrome (HUS), a systemic condition characterised by microangiopathic haemolytic anaemia, thrombocytopenia, and acute kidney injury, which can result death (28). High risk groups for HUS development following STEC infection are young children under five years of age and the elderly (29). Severity of disease, particularly HUS development, is linked to the presence of specific *stx* subtypes (30–32). Of the known subtypes *stx2a* has been associated with the most severe clinical disease outcomes regardless of associated serogroup (23,33).

STEC is a notifiable disease in England and isolates have undergone whole genome sequencing as part of routine surveillance and outbreak investigation by the United Kingdom Health Security Agency (UKHSA) since 2016 (34,35). Leveraging this substantial genomic dataset, we aimed to investigate the temporal dynamics of clinically severe *stx2a* and assess the acquisition of this subtype across O serogroups in England.

## Methodology

### Data download and initial data filtering

Clinically relevant samples from patients with suspected STEC infection underwent whole genome sequencing at UKHSA as part of national surveillance and outbreak investigation, with typing and confirmation processes performed as previously described (34). As part of this, basic metadata including collection year and country of sequencing were deposited within EnteroBase alongside genomic assemblies.

An initial search was performed within the “*Escherichia/Shigella”* EnteroBase database to identify isolates uploaded by UKHSA. Relevant location information was configured to only include isolates from England (per UKHSA remit) and filtering of produced results to only include those submitted by UKHSA, identified through relevant entries within the ‘lab contact’ metric. Non-*E. coli* and *Shigella* species were identified and removed from the results using the ‘species’ metric. Samples of human origin were chosen where consensus of metadata matched across ‘source niche’, ‘source type’ and ‘source detail’ metrics, with samples of non-human origin removed. Data were further restricted to only include isolates sequenced between 2016 and 2024. Samples were removed where raw read sequencing data was unavailable. For the remaining entries, genomic assemblies and associated metadata were downloaded on 13/01/2025. All accession numbers for sequences included in this study are available in **Supplementary Table 1**. Filtering was undertaken both within EnteroBase and using the dplyr (v.1.1.4) package in R (v4.4.3) within RStudio (v2024.12.1+563) (36–38).

### Serotyping, *stx* gene information and deduplication

Serotype and *stx* gene presence were determined as part of UKHSA pipelines as previously described (34,39–41). All serotype and *stx* gene information for sequences included in this study are available in **Supplementary Table 1**. UKHSA detects *stx1a, stx1c, stx1d* and *stx1e* alongside *stx2a-g* as part of routine surveillance. Deduplication was carried out to remove repeat sampling, apart from in the case of mixed infection (different reported STEC serotypes from the same patient) using dplyr. Internal quality control isolates were further removed. Data with a non-standard or no O or H serogroup listed (e.g. O123O186 or O11SSI) were removed. Overall, this gave a final dataset containing 19,429 *E. coli* genomes, which had serotyping information from O and H antigens, alongside *stx* gene information, where applicable. The *stx* genes present within each isolate were referred to as the *stx* profile. Where more than one gene was present per isolate, these were listed in both numerical and alphabetical order, denoted in quotation marks – i.e. “*stx2a, stx2c”*. A presence absence matrix was created for the given *stx* profile using dplyr, separating subtypes per column and introducing a “-” column for isolates with no *stx* gene. All isolates containing *stx* and *stx2a* were filtered for using this matrix.

### STEC and *stx2a* sub-dataset quantification

The janitor (v2.2.1) package in R was used to tidy column names (42). Data counts (e.g. reported isolates by collection month, O antigen presence, *stx* presence) and filtering were undertaken using group_by() and count() within dplyr. All graphical analysis was carried out in ggplot2 (v3.5.1), with the scales (v1.3.0) package used to support visualisation and patchwork (v1.3.0) used to panel graphics together (43–45).

### Acquisition of *stx2a* and *stx2a-*containing *stx* profiles

The first reported date of *stx2a* alone or within further *stx* profiles and per O serogroup was calculated using the first reported collection date and month of an isolate using dplyr. The first reported date was also referred to as the acquisition date, with dates reflective of the first report within the study period, and not reflective of historic acquisition. These were visualised graphically using ggplot2.

## Results

### UK STEC Overview

19,429 *E. coli* isolates of human origin were present in the dataset, reported between 2016 and 2024. Of these, 66.3% (n = 12,888) of isolates were STEC, harbouring up to four stx genes. On average, 1432 STEC isolates were recorded within our dataset each year (SD = 562.5, SE = 187.51) (**Figure 1A**). Seasonality was observed, with a higher number of isolates seen within the summer and early autumn months each year.

**Figure 1.**
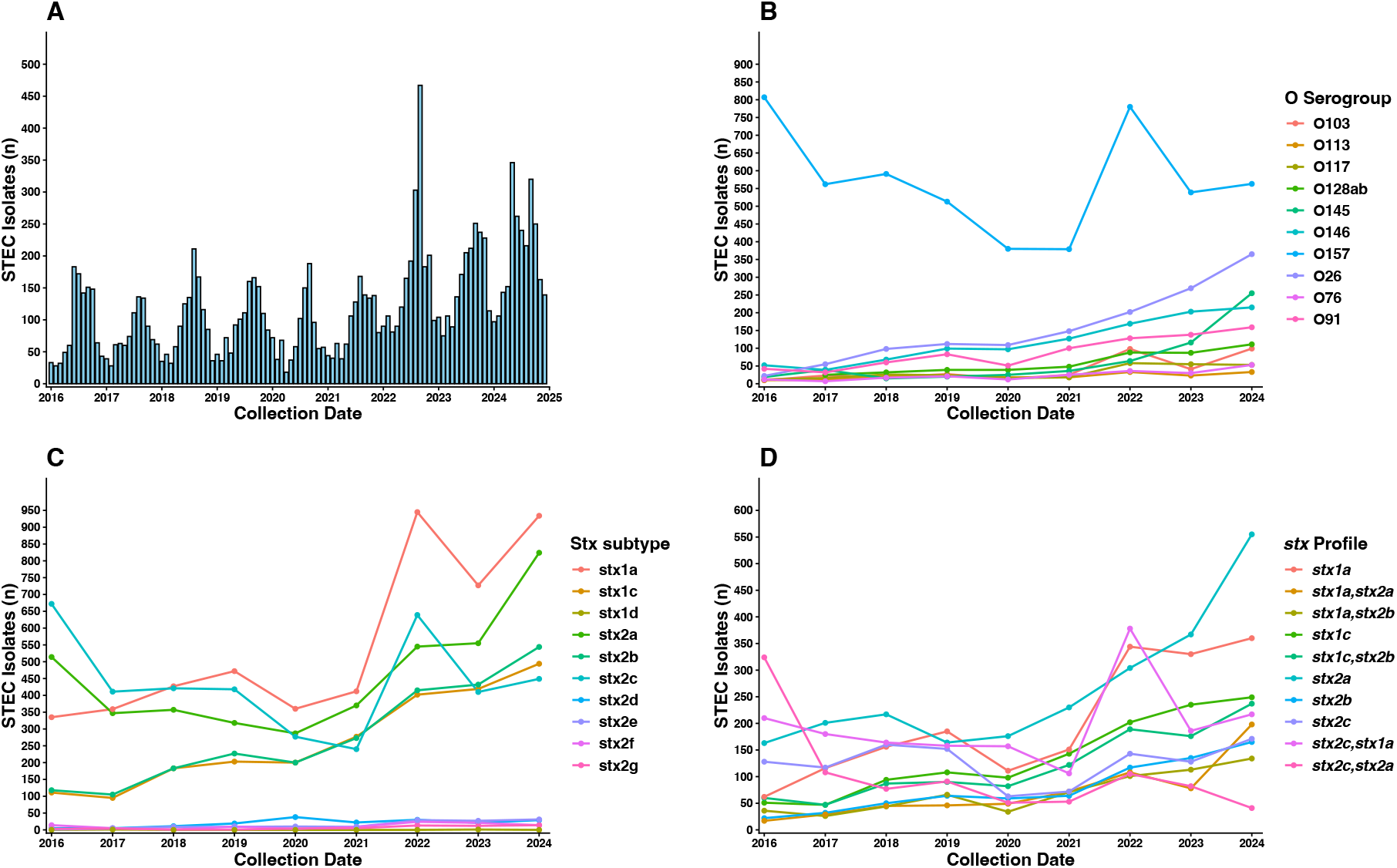
Prevalence and diversity of STEC isolates reported in England, 2016 - 2024. **A** – Report date of STEC isolates across the study period. **B** – Yearly distribution of the 10 most frequently observed O serogroups amongst STEC isolates. 136 unique O serogroups identified overall, remaining serogroups detected at low frequencies. **C** – Yearly distribution of stx subtypes identified within STEC isolates. **D** – Yearly distribution of the 10 most frequently observed stx profiles. 44 unique profiles identified overall, remaining profiles were detected at low frequencies. All isolates originate from human samples reported to UKHSA in England between 2016 and 2024.

### STEC diversity

136 O serogroups and 314 OH serotypes were represented across STEC isolates. The most frequently represented serogroup was STEC O157 (39.7%, n = 5114), however most isolates belonged to non-O157 serogroups (60.3%, n = 7774). Three of the “Big Six” non-O157 STEC serogroups were present among the top 10 reported O serogroups – O26 (10.7%, n = 1380), O145 (4.6%, n = 589) and O103 (2.8%, n = 356). The remaining three were present in the dataset but at lower levels (O45 – 0.2%, n = 26, O111 – 1.1%, n = 141, O121 – 0.1%, n = 10). Despite most isolates belonging to non-O157 serogroups, the O157 serogroup – particularly represented by the O157:H7 serotype – was the most commonly reported serogroup each year. However, non-O157 serogroups continued to rise in prevalence each year (**Figure 1B**).

### *stx* gene presence

10 *stx* gene subtypes were recorded across STEC isolates (**Table 1**), with a total of 18,375 *stx* genes, accounting for multiple genes present within some isolates. The most frequently detected *stx* subtypes were *stx1a* (38.6%, n = 4,971), *stx2a* (31.9%, n = 4,117) and *stx2c* (30.6%, n = 3,937).

**Table 1.** *stx* gene subtype presence across STEC isolates reported in England, 2016 – 2024.

| *stx* subtype | STEC Isolates (n) | STEC Isolates (%) |
| --- | --- | --- |
| *stx1a* | 4971 | 38.6 |
| *stx2a* | 4117 | 31.9 |
| *stx2c* | 3937 | 30.6 |
| *stx2b* | 2497 | 19.4 |
| *stx1c* | 2384 | 18.5 |
| *stx2d* | 180 | 1.4 |
| *stx2e* | 129 | 1.0 |
| *stx2f* | 101 | 0.8 |
| *stx2g* | 57 | 0.4 |
| *stx1d* | 2 | <0.1 |

*Recorded stx gene presence across 12,888 STEC isolates originating from human samples reported to UKHSA in England between 2016 and 2024. A total of 18,375 genes reported across STEC isolates, accounting for those harbouring more than one stx gene.*

Five of the ten subtypes were consistently found in over ∼100 reported isolates per year – *stx1a, stx1c, stx2a, stx2c* and *stx2b*. These five subtypes represented 97.4% of all *stx* genes. The remaining five *stx* subtypes appeared in low levels and were not present in more than 50 reported isolates each year (**Figure 1C**).

### *stx* profile prevalence

The *stx* genes present within each isolate were defined as its *stx* profile, denoted by inverted commas – e.g. *“stx1a, stx2a” or “stx2a”*. 44 unique *stx* profiles were reported across STEC isolates, with each profile containing either one, or a combination of up to four *stx* genes (**Table 2**). All ten *stx* subtypes were present individually, representing ten *stx* profiles. The remaining 34 *stx* profiles contained more than one *stx* gene – 18 contained two genes, 15 contained three and one contained four genes. All reported *stx* subtypes appeared across these remaining profiles except for *stx1d* and *stx2f* which only appeared in isolation.

**Table 2.** *stx* profile presence across STEC isolates reported in England, 2016 – 2024.

| *stx* profile | STEC Isolates (n) | STEC Isolates (%) |
| --- | --- | --- |
| "*stx2a*" | 2377 | 18.4 |
| "*stx1a*" | 1815 | 14.1 |
| "*stx1a, stx2c*" | 1756 | 13.6 |
| "*stx1c*" | 1227 | 9.5 |
| "*stx2c*" | 1134 | 8.8 |
| "*stx1c, stx2b*" | 1090 | 8.5 |
| "*stx2a, stx2c*" | 932 | 7.2 |
| "*stx2b*" | 708 | 5.5 |
| "*stx1a, stx2a*" | 642 | 5.0 |
| "*stx1a, stx2b*" | 625 | 4.9 |
| "*stx2f*" | 101 | 0.8 |
| "*stx2d*" | 81 | 0.6 |
| "*stx1a, stx2a, stx2c*" | 81 | 0.6 |
| "*stx2e*" | 73 | 0.6 |
| "*stx2g*" | 53 | 0.4 |
| "*stx2a, stx2e*" | 46 | 0.4 |
| "*stx1c, stx2b, stx2d*" | 36 | 0.3 |
| "*stx2b, stx2d*" | 10 | 0.1 |
| "*stx1a, stx2a, stx2e*" | 10 | 0.1 |
| "*stx1c, stx2d*" | 9 | 0.1 |
| "*stx2c, stx2d*" | 9 | 0.1 |
| "*stx1a, stx2d*" | 7 | 0.1 |
| "*stx1a, stx2b, stx2d*" | 7 | 0.1 |
| "*stx1a, stx2a, stx2d*" | 6 | 0.1 |
| "*stx1a, stx1c*" | 5 | <0.1 |
| "*stx1a, stx1c, stx2b*" | 5 | <0.1 |
| "*stx2b, stx2c*" | 5 | <0.1 |
| "*stx1a, stx2c, stx2d*" | 5 | <0.1 |
| "*stx1a, stx2a, stx2c, stx2d*" | 4 | <0.1 |
| "*stx2a, stx2g*" | 3 | <0.1 |
| "*stx2a, stx2b*" | 3 | <0.1 |
| "*stx2a, stx2d*" | 3 | <0.1 |
| "*stx1c, stx2a*" | 3 | <0.1 |
| "*stx1c, stx2b, stx2c*" | 3 | <0.1 |
| "*stx1d*" | 2 | <0.1 |
| "*stx1c, stx2a, stx2b*" | 2 | <0.1 |
| "*stx2a, stx2c, stx2d*" | 2 | <0.1 |
| "*stx1c, stx2c*" | 2 | <0.1 |
| "*stx1a, stx2g*" | 1 | <0.1 |
| "*stx1a, stx2a, stx2b*" | 1 | <0.1 |
| "*stx2b, stx2c, stx2d*" | 1 | <0.1 |
| "*stx2a, stx2b, stx2c*" | 1 | <0.1 |
| "*stx1c, stx2a, stx2c*" | 1 | <0.1 |
| "*stx1a, stx1c, stx2c*" | 1 | <0.1 |

*stx* profile presence across 12,888 STEC isolates originating from human samples reported to UKHSA in England between 2016 and 2024. "*stx* profile" defined as all *stx* genes present within each isolate. 6,541 isolates of the original 19,429 *E. coli* isolates used in the study were *stx* negative and therefore not shown within the table.

“*stx2a”* was the most observed *stx* profile, present in 18.44% (n = 2377) of STEC isolates, this was followed by *“stx1a”* present in 14.08% of STEC isolates (n = 1815) and *“stx2c, stx1a”* present in 13.63% (n = 1756) of STEC isolates. *stx2a* was the most distributed subtype, present in sixteen further *stx* profiles. This was followed by *stx1a* and *stx2c* in 15 and 14 profiles respectively.

Displaying the number of reported isolates across the top ten *stx* profiles per year (**Figure 1D**) revealed an overall rise post-2020, aside from *“stx2a, stx2c”*, and the steepest overall rise in profiles *“stx2a”* and “*stx1a”*. The remaining 34 *stx* profiles were present in low numbers, under ∼25 isolates per year each. *“stx2a”* saw the biggest increase of all profiles post-2020.

### *stx2a* presence and fluctuation

Given the rise in frequency observed across isolates with the *stx2a* subtype and *stx* profiles containing *stx2a*, alongside its association with clinical severity, we sought to identify further trends in prevalence and acquisition of *stx2a* within the UK. A large rise was seen in the number of isolates harbouring *stx2a* post 2020, jumping from 287 in 2020 and 824 in 2024, with 318-514 isolates reported each year in between (**Figure 2A**). Seasonal fluctuation was observed, with more *stx2a*-containing isolates reported in the summer and autumn months each year. Of the 17 *stx* profiles containing *stx2a*, only three were substantially reported across the nine-year period – *“stx2a”* (n=2377), *“stx2a, stx2c”* (n = 932) and *“stx1a, stx2a”* (n = 642) (**Figure 2B**). The remaining profiles containing *stx2a* were present in low levels.

**Figure 2.**
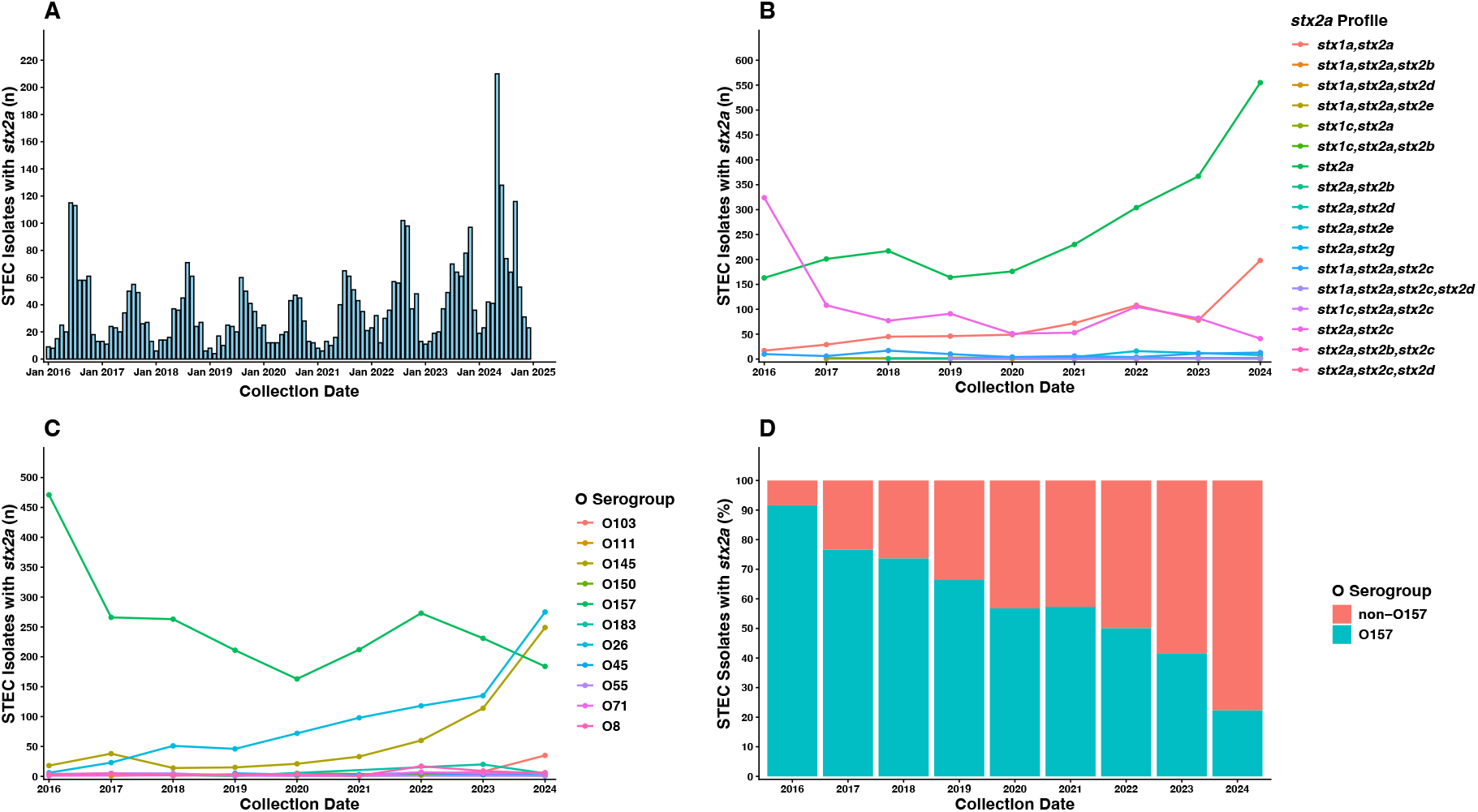
Prevalence and diversity of stx2a in England, 2016 - 2024. **A** – Report date of STEC isolates harbouring stx2a across the study period. **B** – Yearly distribution of stx profiles amongst stx2a harbouring STEC isolates. **C** – Yearly distribution of the 10 most frequently observed O serogroups amongst stx2a harbouring STEC isolates. 78 unique O serogroups identified overall, remaining serogroups detected at low frequencies. **D** – Proportional distribution of O157 vs non-O157 serogroups among stx2a harbouring isolates. All isolates originate from human samples reported to UKHSA in England between 2016 and 2024.

### *stx2a* distribution amongst serogroups

*stx2a* was found within 78 unique O serogroups, with the highest representation in O157 (55.2%, n = 2274), O26 (20.0%, n=824) and O145 (13.7%, n=562). Despite a strong presence from the O157 serogroup, 44.8% (n = 1843) of isolates harbouring *stx2a* were from 77 unique non-O157 serogroups. The number of O157 isolates containing *stx2a* remained relatively stable throughout the study period but the number of non-O157 isolates containing *stx2a* steadily increased, with a significant rise each year post-2021 (**Figure 2C)**. Proportionately, the number of *stx2a* harbouring O157 isolates fell steadily from 2016, mirrored by an increase of non-O157 serotypes, with more diversity seen in these groups across the study period (**Figure 2D**).

Three main serogroups were responsible for the high levels of *stx2a* – O157, O26 and O145 – specifically STEC serotypes O157:H7, O26:H11 and O145:H28 (**Figure 3A**). Both STEC O26:H11 and STEC O145:H8 saw a steady overall increase in reported isolates between 2016 and 2023, ranging from 18-113 (STEC O145:H28) and 6-135 (STEC O26:H11) each year. The greatest increase for both serotypes was in 2024, with 249 reported isolates for STEC O145:H28 (+136 from 2023) and 274 for STEC O26:H11 (+139 from 2023). In our dataset, STEC O157:H7 cases decreased across the study period, falling from 471 isolates in 2016 to 184 in 2024. Non-O157 groups O145 and O26 were reported more frequently than O157 in 2024.

**Figure 3.**
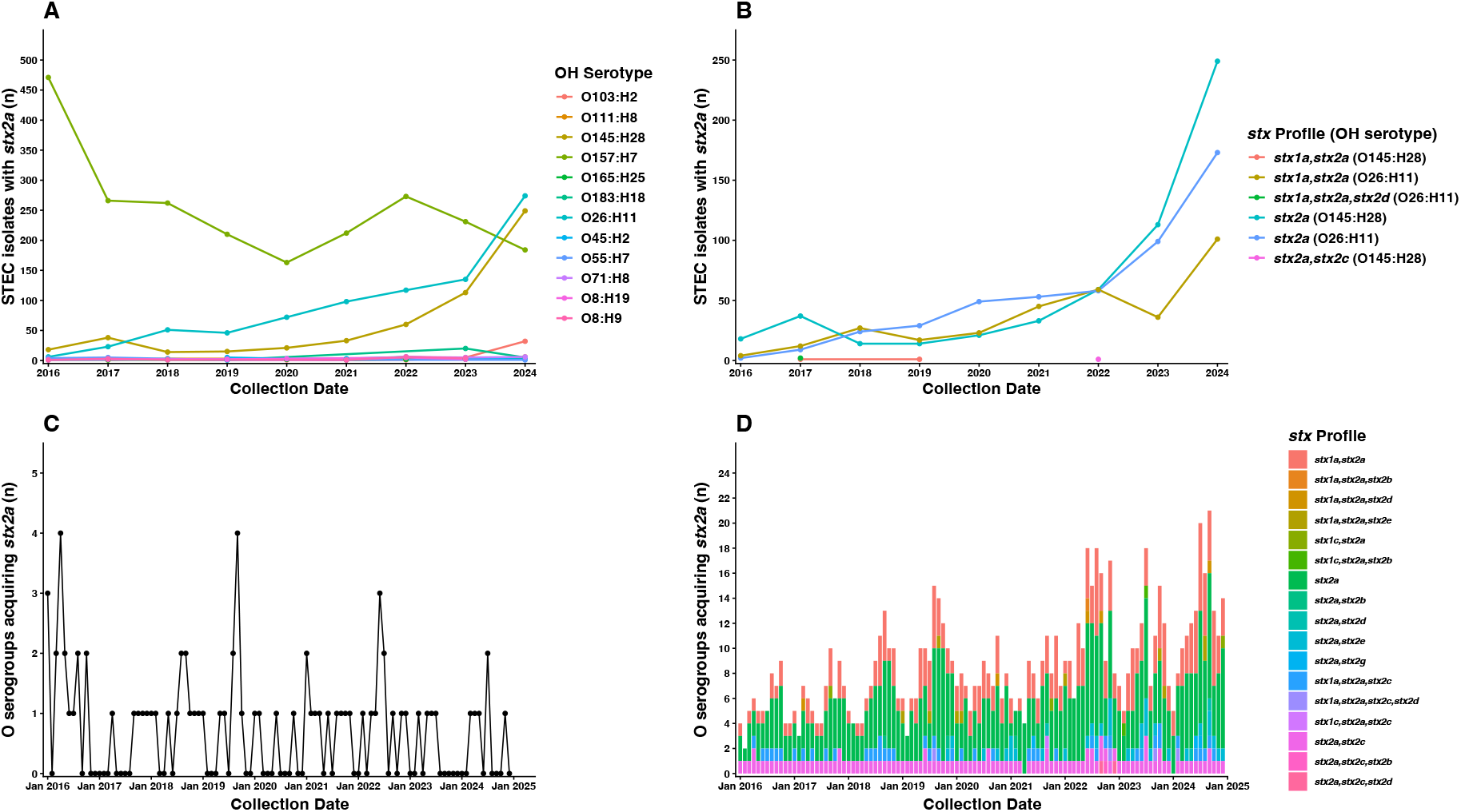
Drivers of increase and acquisition of stx2a isolates in England, 2016 - 2024. **A** – Yearly distribution of the top 10 O serotypes reported per year across stx2a-harbouring STEC isolates. **B** – Yearly distribution of stx2a-containing profiles among STEC O26:H11 and STEC. O145:H28. **C** – Monthly distribution of O serogroups acquiring the stx2a subtype. **D** – Monthly distribution of O serogroups acquiring stx2a as part of any stx2a-containing stx profile. All isolates originate from human samples reported to UKHSA in England between 2016 and 2024.

STEC O26:H11 was represented by three *stx2a* profiles – *“stx2a”, “stx1a, stx2a” and “stx1a, stx2a, stx2d’*. However, only STEC O26:H11 STEC with profiles *“stx2a”* and *“stx1a, stx2a”* contributed to the increase in the serotype, with *“stx2a”* displaying with the greatest overall increase. STEC O145:H28 was also represented across three *stx2a* profiles – *“stx2a”, “stx1a, stx2a”* and *“stx2a, stx2c”. “stx2a”* was the sole driver of the increase, with remaining isolates appearing minimally (max 2 isolates in sporadic years) (**Figure 3B**).

### *stx2a* acquisition

Within our genomic dataset, isolates with *stx2a* were reported from the start of the study period in January 2016. Of the 17 *stx* profiles containing *stx2a*, variation was seen in the first recorded appearance (acquisition date) of each (**Table 3**). The dates of these spanned evenly across the study period, representing steadily increasing acquisition of and diversity within isolates harbouring *stx2a*.

**Table 3.** First reported dates stx gene profiles harbouring *stx2a*.

| stx Profile | First Recorded Appearance (Date) |
| --- | --- |
| "stx1a, stx2a" | January 2016 |
| "stx2a" | January 2016 |
| "stx2a, stx2c" | January 2016 |
| "stx1a, stx2a, stx2c" | April 2016 |
| "stx1a, stx2a, stx2d" | March 2017 |
| "stx1c, stx2a" | September 2017 |
| "stx2a, stx2b" | September 2018 |
| "stx1a, stx2a, stx2e" | January 2019 |
| "stx1a, stx2a, stx2c, stx2d" | August 2019 |
| "stx2a, stx2e" | November 2019 |
| "stx2a, stx2g" | June 2021 |
| "stx2a, stx2d" | September 2021 |
| "stx1c, stx2a, stx2c" | September 2021 |
| "stx1a, stx2a, stx2b" | June 2022 |
| "stx2a, stx2c, stx2d" | September 2022 |
| "stx2a, stx2b, stx2c" | October 2022 |
| "stx1c, stx2a, stx2b" | February 2023 |

First recorded appearance of each stx profile containing the stx2a gene within the dataset. An "stx profile" is defined as the combination of stx genes present within an individual isolate. The first recorded appearance date is referred to as the acquisition date. UKHSA began whole genome sequencing in 2016, with acquisition dates reflecting the earliest detection within our dataset and do not necessarily represent the timing of emergence within the wider population.

The diversity of O serogroups harbouring and acquiring *stx2a* also increased over time. The number of new O serogroups acquiring the *stx2a* subtype varied between 0-4 per month across the study period. In 2016, there were 17 O serogroups which harboured *stx2a*, with 5-11 new O serogroups acquiring *stx2a* each year between 2017 to 2024 (**Figure 3C**). Post-2020, O serogroups harbouring *stx2a* increased from four in 2020 to 10 and 11 in 2021 and 2022 respectively.

The highest number of O serogroups acquiring *stx2a* within any *stx2a*-containing profile in any month was 21 in September 2024. When broken down by specific *stx* profile, three profiles are responsible for driving this acquisition – “*stx2a*”, “*st2a, stx2c*” and “*stx1a, stx2a*” (**Figure 3D**). These were also the highest reported profiles containing *stx2a* per year.

## Discussion

Here, we have described the temporal dynamics and acquisition of Shiga toxin subtype *stx2a* found across STEC in England. *stx2a* was present in just under a third of all STEC isolates in our dataset, highlighting a substantial portion of notified STEC cases had the potential to cause severe disease and progression to HUS. By characterising STEC serotype and *stx* subtype across a 9-year period from public health surveillance data, we have observed an increasing trend in STEC isolates harbouring *stx2a* and quantified the diversification of *stx2a* across *E. coli* serogroups and presence alongside other *stx* subtypes.

The number of isolates harbouring *stx2a* rose each year throughout the study period, particularly post-2020. Although STEC O157 carrying *stx2a* continue to be of concern to public health, notifications of these isolates have decreased in recent years. Whilst being represented across 78 serogroups, two serotypes were largely responsible for driving the increase in detection of *stx2a* – STEC O26:H11 and STEC O145:H28. Previous assessment of non-O157 STEC serogroups in England saw an association of both of these serotypes with greater risk of severe disease, alongside higher hospitalisation rates and reported HUS (23). The increase of *stx2a* caused by STEC O145:H28 was driven by isolates harbouring the “*stx2a*” *stx* profile only, however, isolates of STEC O26:H11 harboured either “*stx2a*” in isolation or “*stx1a, stx2a*”, highlighting two *stx2a* profiles of concern of this serotype. *stx1a* is commonly found alongside *stx* subtypes and is known to cause mild disease (20). STEC O26:H11 and STEC O145:H28 encoding *stx2a* were present in the dataset from the start of the study period in 2016. Therefore, it is likely these strains were circulating in the population prior to this date. Further investigation into the genomic background of STEC O145:H28 and STEC O26:H11 isolates with the profiles described may reveal additional factors, such as virulence genes, genes facilitating persistence and survival in the environment, and/or enhancing transmission between animals and people, which may be contributing to the observed increase.

These isolates were distributed across a wide range of serogroups in the STEC dataset for England, with the number of serogroups acquiring *stx2a* increasing each month, displaying temporal diversification across the E. coli population structure. It remains of concern that *stx2a* is becoming more widespread, giving rise to a wider number of different serotypes with the potential to cause severe disease. Of the *stx* profiles, isolates harbouring “*stx2a*” present on its own were the main driver of the yearly increase of STEC harbouring *stx2a. stx2a* also was present within 17 different *stx* profiles – the highest for any recorded stx subtype. Our study shows the value of including tracking STEC by *stx* subtype. The continual inclusion of this in routine public health surveillance may ultimately help identify emerging highly pathogenic serotypes associated with causing progression to HUS, improving diagnosis and allowing for earlier clinical intervention.

*stx2a* is not the only *stx* subtype capable of causing HUS, but it is associated with the highest progression to severe disease, highlighting this subtype as a public health priority. Further subtypes including *stx2c, stx2d* and *stx2f* have further been known to cause disease progression to HUS, however in fewer cases in comparison (23). By continuing to place emphasis on *stx* subtyping as part of routine surveillance, these further subtypes with increased disease severity will be continued to be identified, allowing effective downstream measures to be taken. The ten represented *stx* in this study (*stx1a, stx1d, stx1c, stx1d, stx2a-f*) are present as part of routine surveillance by UKHSA (34). The effects of novel subtypes of Stx2 (*stx2g-o)* on disease severity in humans largely remain unknown, but future inclusion of these within routine surveillance may further contribute to successful public health responses towards STEC.

Over the last 15 years, there has been a shift in STEC diagnostics, with PCR detection of the stx gene replacing traditional culture-based approaches. As methods for molecular serotyping were largely geared towards the O157 serogroup, particularly the O157:H7 serotype, non-O157 groups were historically underreported. The shift to PCR detection seen a subsequent increase in non-O157 serogroup notifications, which is a contributing factor to increasing trends. However, the evidence we present here suggests the increasing trend is not solely the result of reporting bias due to the implementation of PCR.

## Conclusion

STEC isolates harbouring *stx2a* have both increased and diversified over the last decade in England. *stx2a* – linked with more severe disease outcomes, including HUS – was present in a substantial portion of isolates reported to UKHSA since 2016 and displayed a year-on-year increase. Three subgroups of differing serotypes and/or *stx2a* profiles – STEC O26:H11 with “*stx2a”*, STEC O26:H11 with “*stx1a, stx2a*” and STEC O145:H28 “*stx2a*” – were responsible for driving the increase, posing a risk to public health that should be monitored closely. Further investigation into the reservoirs and transmission pathways of these subgroups may strengthen evidence for informing food safety policy and public health guidance.

## Supporting information

Supplementary Table 1. Related Accession Numbers

## Ethical approval and consent to participate

The authors declare that there was no requirement for ethical approval for this submission. This work was undertaken to inform the delivery of patient care and to prevent the spread of infection, defined as USUAL PRACTICE in public health and health protection.

## Author contributions

**EHH** – conceptualisation, formal analysis, visualisation, writing – original draft, writing – review & editing. **EVR** – conceptualisation, resources, supervision, writing – review & editing. **DRG** – conceptualisation, resources, supervision, writing – review & editing. **CJ** – conceptualisation, resources, supervision, writing – review & editing. **GCL** – conceptualisation, supervision, writing – review & editing. All authors agreed on the final version of the manuscript.

## Acknowledgements

The authors would like to extend their gratitude to the wider Gastrointestinal Bacteria Reference Unit (GBRU) within UKHSA for creation of and facilitating access to data.

